# The lncRNA HOTAIR/EZH2 interaction inhibitor AC1Q3QWB (AQB) attenuates fibrotic SSc skin tissue re-modelling

**DOI:** 10.64898/2026.06.12.730368

**Authors:** Christopher W. Wasson, Panji Mulipa, Sophie L. Dibb, Esther Perez Barreiro, Rebecca. L. Ross, Francesco Del Galdo, Natalia. A. Riobo-Del Galdo

## Abstract

**Objectives:** The long non-coding RNA HOTAIR has previously been shown to drive pro-fibrotic gene expression in SSc dermal fibroblasts through its ability to interact with EZH2. Targeting EZH2 enzymatic activity can reverse HOTAIR mediated pro-fibrotic gene expression but its many functions make it an undesirable therapeutic target for SSc. Recently inhibitors selectively targeting the HOTAIR/EZH2 interaction have been developed. The aim of this study was to characterise the ability of one of these inhibitors to modulate SSc tissue remodelling.

**Methods:** Explanted healthy and SSc dermal fibroblasts were treated with the HOTAIR/EZH2 inhibitor AC1Q3QWB (AQB) (20µM) for 48 hours. In addition, healthy dermal fibroblasts were transduced with a lentivirus encoding HOTAIR or a scrambled control. Conditioned media from healthy, SSc and HOTAIR-expressing dermal fibroblasts was used to stimulate human keratinocytes (HaCaTs). Scramble control and HOTAIR expressing fibroblasts were grown in 3D skin equivalents containing primary keratinocytes and keratin 9 (K9) immunohistochemistry performed.

**Results:** AQB inhibits pro-fibrotic gene expression in HOTAIR expressing dermal fibroblasts, validating the specificity of the inhibitor. In SSc patient dermal fibroblasts, AQB blocked pro-fibrotic gene expression but did not affect gene expression in healthy dermal fibroblasts. SSc patient skin was shown to express high levels of the palmoplantar specific K9 and Epithelial to Mesenchymal transition (EMT) markers. Through co-culture experiments we showed these effects were mediated by SSc dermal fibroblasts. This tissue remodelling was disrupted when HOTAIR/EZH2 interaction was inhibited in the fibroblasts with AQB.

**Conclusions:** We have shown for the first time that directly inhibiting HOTAIR/EZH2 interaction blocks pro-fibrotic gene expression in SSc fibroblasts and tissue re-modelling found in SSc patient skin. This may represent a novel therapeutic intervention.

## Introduction

The long non-coding RNA HOTAIR has previously been shown to be a master regulator of chromatin modification (1). The 2158 nucleotides-long HOTAIR adopts a specific fold that serves as a protein-binding scaffold. HOTAIR is known to recruit the Polycomb repressor Complex (PRC2) and the REST/co-REST complex to promoters leading to regulation of gene expression by methylation and demethylation, respectively (1). HOTAIR interacts with the enzymatic subunit of the PRC2, Enhancer of Zeste homolog 2 (EZH2), through its 5’ domain and with REST/coREST through a 3’ domain (2). Several studies have demonstrated that the scaffolding function of HOTAIR directs the epigenetic remodelling complexes to specific genomic loci in a cell type and disease-specific manner (1, 2).

Previous work from our group has shown HOTAIR is highly expressed in the Systemic Sclerosis (SSc) patient skin and in isolated primary and immortalised dermal fibroblasts (3). Overexpression of HOTAIR induces pro-fibrotic gene expression in dermal fibroblasts from healthy controls, in part through co-operation with the PRC2 complex (4, 5). Inhibition of the EZH2 blocks pro-fibrotic gene expression in SSc fibroblasts and reverses the ability of HOTAIR to induce pro-fibrotic gene expression in healthy fibroblasts (3, 5). In addition, HOTAIR can activate the Notch signalling pathway in an EZH2-dependent manner, which in turn drives pro-fibrotic gene expression (3,4). Further analysis revealed the co-REST complex played an important role in SSc myofibroblasts activation when the cells were stimulated with pro-fibrotic stimuli, but this role was largely independent of HOTAIR (6). This evidence suggests that targeting the EZH2-dependent effects of HOTAIR could have therapeutic value in SSc.

Several compounds that specifically target EZH2 are undergoing clinical trials for a range of cancers (7). Unfortunately, the side effects associated with these compounds limit their scope for clinical development in SSc. In this context, specific therapeutic options that target the HOTAIR-PRC2 interphase would be more suitable for development as antifibrotics. Recently, through leveraging in silico modelling through MC-Fold and MC Sym programmes, a series of compounds specifically targeting HOTAIR interaction with EZH2 have been developed (8, 9). Through a series of RIP and ChIRP assays, these compounds were shown to selectively interfere with the HOTAIR/EZH2 interaction (10).

Here we tested the biological activity of one these compounds, AC1Q3QWB (hereafter referenced as AQB) to suppress the pro-fibrotic effects of the HOTAIR-EZH2 complex without interfering with the other functions of EZH2. Our findings reveal that AQB is a potent and specific HOTAIR/EZH2 inhibitor that reduces profibrotic gene expression and tissue remodelling in SSc.

## Materials and Methods

### Patient cell lines

Full thickness skin biopsies were surgically obtained from the forearms of four adult healthy controls and four adult patients with recent onset SSc, defined as a disease duration of less than 18 months from the appearance of clinically detectable skin induration. All patients satisfied the 2013 ACR/EULAR criteria for the classification of SSc as defined by LeRoy et al (11). All participants provided written informed consent to participate in the study. Informed consent procedures were approved by NRES-011NE to FDG. Fibroblasts were isolated and established as previously described (12, 13). Primary cells were immortalized using human telomerase reverse transcriptase (hTERT) to produce healthy control hTERT and SSc hTERT.

### Cell culture

hTERT patient fibroblasts were maintained in Dulbecco’s modified Eagle medium (DMEM) (Gibco) supplemented with 10% FBS (Sigma) and penicillin-streptomycin (Sigma). Fibroblasts were treated with AQB (5-30μM) for 48 hours in a humidified incubator at 37C and 5% CO2. GSK-126 (an EZH2 methylation transferase inhibitor) was used at a final concentration of 5μM (LKT Laboratories:G7340). FH535 (a β-Catenin inhibitor) was used at a final concentration of 10μM (Sellekchem) Healthy dermal fibroblasts were serum starved for 24 h in DMEM containing 0.5% FBS and stimulated with 10 ng/ml TGF-β (R&D systems).

### Lentiviral Transduction

Fibroblasts were grown from healthy control forearm biopsies and immortalised using retrovirus expressing human telomerase (hTERT) as previously outlined (12, 13). HOTAIR expression was then induced by transduction with GIPZ lentiviruses carrying HOTAIR gene sequence or scrambled RNA sequence as control in frame with puromycin resistance gene and GFP fluorochrome gene. (Open Biosystems, Surrey, UK). For this purpose, fibroblasts were seeded at 50% confluence and infected with lentiviral particles in serum free DMEM and incubated for 6h, after which an additional 1ml of DMEM containing 10% FCS was added and the cells were incubated for a further 72h. Stably transduced fibroblasts were positively sorted for GFP fluorescence employing Fluorescence Activated Cell Sorting in sterile conditions (BD INFLUX). Positively sorted cells were further selected in media containing 1.0μg/ml puromycin (Life Technologies) for 10 days.

### 3D skin equivalents

Collagen matrix solution containing dermal fibroblasts was seeded into transwell inserts and allowed to solidify. Primary keratinocytes were seeded on top of the collagen matrix. Once at confluence, media was removed from the transwell insert to create an air liquid interface. The skin equivalent was grown at the interface for 8 days at which point it was harvested for histology.

### Western blotting

Total proteins were extracted from fibroblasts in RIPA buffer and resolved by SDS-PAGE (10-15% Tris-Glycine). Proteins were transferred onto Hybond nitrocellulose membranes (Amersham biosciences) and probed with antibodies specific for alpha smooth muscle actin (Abcam ab5694), Notch 1 (Cell signalling 3608), β-catenin (cell signalling 9562), CTGF (abcam ab6992), pSMAD3 (abcam ab40854), Keratin 9 (abcam ab19124), N-Cadherin (Santa Cruz sc59987), Vimentin (Cell signalling 5741) and β-Actin (Sigma A5441). Immunoblots were visualized with species-specific HRP conjugated secondary antibodies (Sigma) and ECL (Thermo/Pierce) on a Biorad chemiDoc imaging system.

### Immunofluorescence

Scramble and HOTAIR expressing fibroblasts were seeded onto coverslips and treated with AQB for 48 hours. The cells were fixed in 4% paraformaldehyde and permeablised with 0.1% trition-x20 for 10 mins. The cells were stained with an alpha-SMA antibody (abcam ab5694) and visualised with a mouse secondary antibody conjugated to alexa-594. Nuclei were visualised by DAPI contained within the mounting media. Scale bars represent 20μm.

### Immunohistochemistry

Immunohistochemistry was performed as previously described (14). Sections were stained with a Keratin 9 (1/200) (abcam ab19124) antibody, visualised using an HRP conjugated mouse secondary and counterstained with haematoxylin.

### Quantitative Real time PCR

RNA was extracted from cells using commercial RNA extraction kits (Zymo Research). RNA (1μg) was reverse transcribed using cDNA synthesis kits (Thermo). QRT-PCRs were performed using SyBr Green PCR kits on a Thermocycler with primers specific for *HOTAIR* (Forward: 5’-GGTAGAAAAAGCAACCACGAAGC; Reverse: 5’-ACATAAACCTCTGTCTGTGAGTGCC), *ACTA2* (Forward: 5’-TGTATGTGGCTATCCAGGCG; Reverse: 5’-AGAGTCCAGCACGATGCCAG), *COL1A2* (Forward: 5’-GATGTTGAACTTGTTGCTGAGC; Reverse: 5’-TCTTTCCCCATTCATTTGTCTT), *CCN2* (Forward: 5’-GTGTGCACTGCCAAAGATGGT; Reverse: 5’-TTGGAAGGACTCACCGCT), *HES1* (Forward: 5’-TACCCAGCCAGTGTCAAC; Reverse: 5’-CAGATGCTGTCTTTGGTTTATCC) and *GAPDH* (Forward: 5’-ACCCACTCCTCCACCTTTGA; Reverse: 5’-CTGTTGCTGTAGCCAAATTCGT). Data were analysed using the ΔΔ Ct method. GAPDH served as a housekeeping gene.

## Results

### AQB attenuates HOTAIR-mediated fibroblast activation

Overexpression of HOTAIR in resting dermal fibroblasts results in myofibroblast activation, including the expression of α-SMA (3). Therefore, as a proof of principle we assessed the ability of the HOTAIR inhibitor AQB (previously described (10)) to modulate pro-fibrotic gene expression in these fibroblasts. Control (Scramble) and HOTAIR-overexpressing fibroblasts were treated with 20µM of AQB for 48 hours. The expression of HOTAIR led to upregulation of α-SMA and CTGF (Figure 1A), consistent with previous reports (3). Treatment with AQB suppressed α-SMA and CTGF proteins levels by 30% and 50%, respectively, (Figure 1A), in HOTAIR-expressing fibroblasts. The inhibitor had minimal effect on the α-SMA expression levels in scramble control fibroblasts (Figure 1A). Moreover, AQB strongly reduced formation of SMA^+^ fibres in HOTAIR-expressing fibroblasts (Figure 1B). In agreement, we observed a strong reduction in transcript levels of *ACTA2* (Figure 1C) and, to a lower extent, of *CCN2* (Figure 1D) in HOTAIR-expressing fibroblasts treated with AQB.

**Figure 1:**
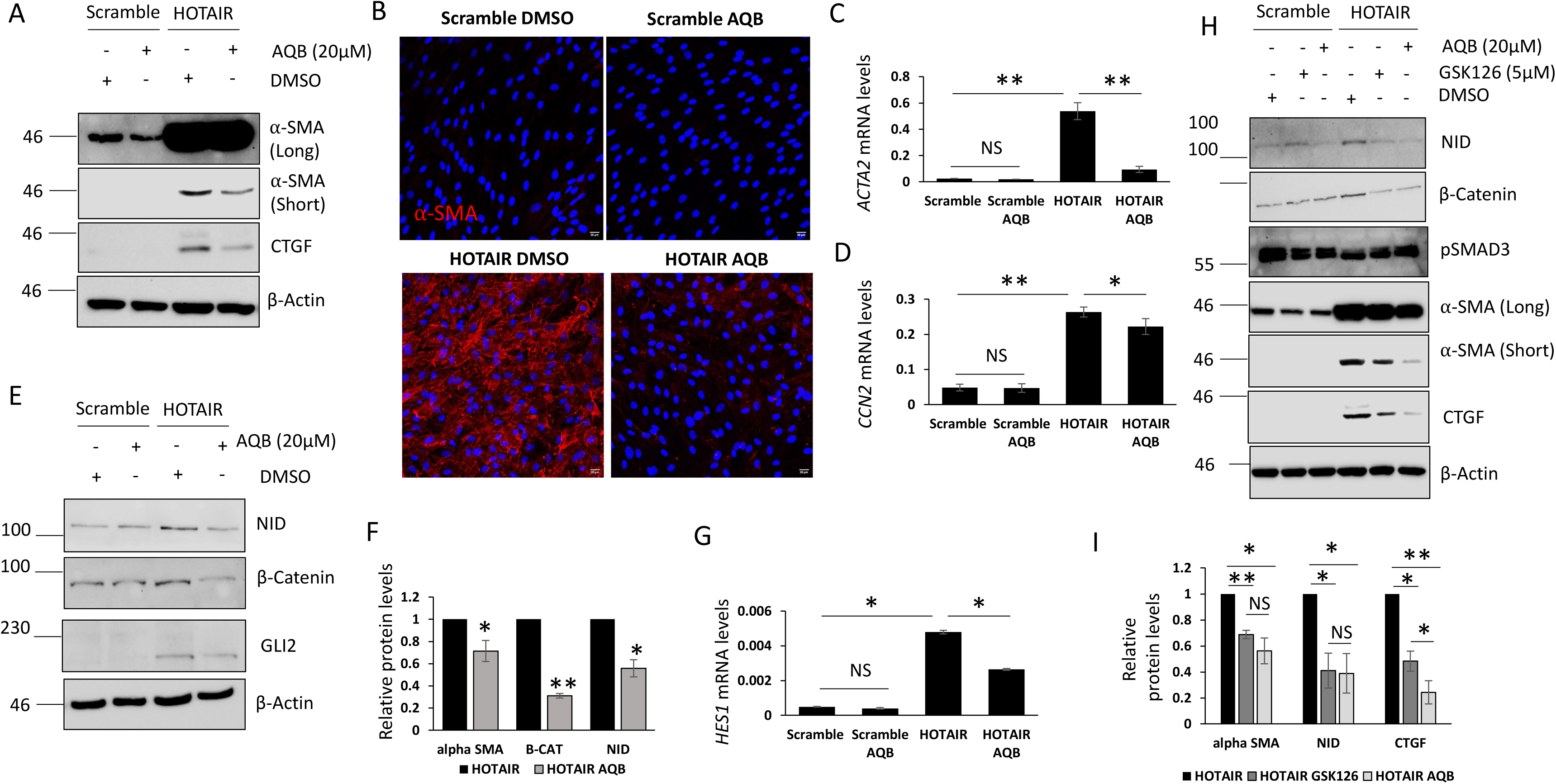
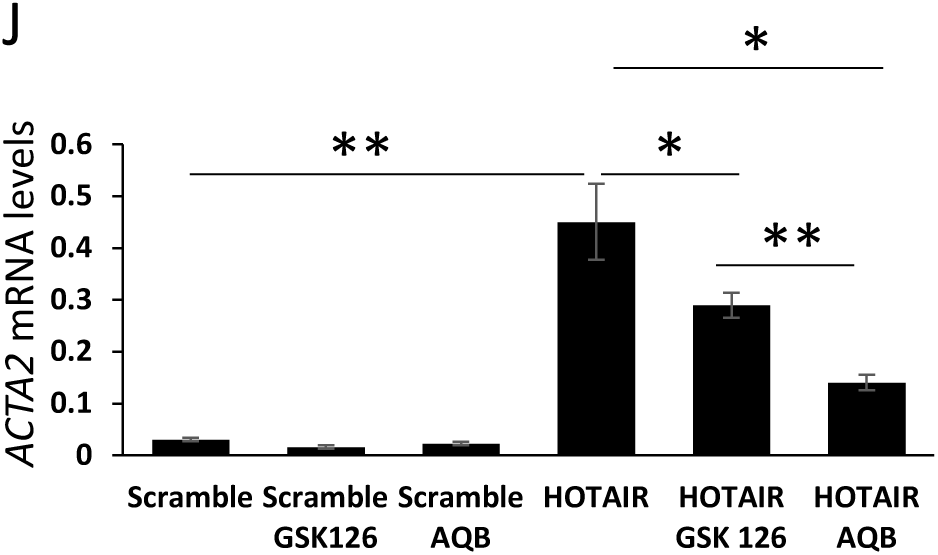
AQB attenuates HOTAIR mediated fibroblast activation. Protein and RNA were extracted from dermal fibroblasts expressing the HOTAIR sequence or a scramble control sequence. In addition, the fibroblasts were treated with the AQB compound (20μM) for 48 hours. (A) α-SMA,, and CTGF, protein levels were assessed by western blot. β-actin was used as a loading control.. α-SMA (B), CCN2 (C) transcript levels were assessed by qPCR. (D) α-SMA staining of scramble and HOTAIR expressing fibroblasts treated with AQB. Staining was visualised with a mouse conjugated alexa594 secondary. (E) NID, GLI2 and β-Catenin protein levels were assessed by western blot. β-actin was used as a loading control. (F) Graph represents densitometry analysis of HOTAIR expressing fibroblasts plus/minus the inhibitors. (G) Hes1 transcript levels were assessed by qPCR. (H) Protein and RNA were extracted from scramble and HOTAIR expressing fibroblasts. In addition, the fibroblasts were treated with the AQB or GSK126 for 48 hours. α-SMA, β-catenin, pSMAD3, CTGF and NID protein levels were assessed by western blot. β-actin was used as a loading control. (I) Graph represents densitometry analysis α-SMA, NID and CTGF protein levels from HOTAIR expressing fibroblasts plus/minus the inhibitors. (G) α-SMA transcript levels were assessed by qPCR.

We previously demonstrated that HOTAIR-mediated fibroblast activation is mediated by activation of the Notch signalling pathway, which in turn induces the expression of the Hedgehog transcription factor GLI2 (4, 6). Activation of the Notch signalling pathway requires cleavage of the Notch receptor to form NID, which acts as a transcriptional co-activator for the pathway. In the same experimental settings, treatment with AQB reduced the level of Notch intracellular domain (NID) (Figure 1E, F). The reduction in NID levels in HOTAIR-expressing fibroblasts treated with AQB suggests the inhibitor efficiently blocked Notch signalling. This was confirmed by a ∼50% reduction of *HES1* transcript levels (a downstream target of the Notch signalling pathway) in cells treated with AQB (Figure 1G). In consequence, GLI2 protein levels were reduced in the HOTAIR expressing fibroblasts treated with AQB (Figure 1E, F),In addition, HOTAIR has been previously shown to increase β-catenin protein levels (15, 16), which have been extensively reported as a central molecular step in SSc myofibroblast activation (15, 17, 18). AQB treatment attenuated the increased β-catenin level in the HOTAIR fibroblasts to levels similar to the scramble control (Figure 1E, F).

Given that TGF-β signalling is a master stimulator of fibroblast activation and that the AQB compound had such as strong inhibitory effect on fibrotic markers expression, we investigated if AQB could inhibit TGFβ/SMAD signalling. Interestingly, neither overexpression of HOTAIR in dermal fibroblasts led to increased SMAD3 activation, indicated by its phosphorylated form pSMAD3, nor this was affected by treatment with AQB (Figure 1H). Taken together this data suggests the HOTAIR/EZH2 interaction inhibitor AQB has the ability to block pro-fibrotic gene expression induced by HOTAIR upregulation in dermal fibroblasts independently of TGF-β/SMAD3 signalling.

Next, we set out to compare the efficacy of AQB compared with enzymatic EZH2 inhibitors. GSK126 is a EZH2 inhibitor currently in clinical development for adjuvant treatment of solid malignancies, and which has previously been shown to block pro-fibrotic gene expression in SSc fibroblasts (3). Both GSK126 and AQB reduced α-SMA, CTGF, NID and β-catenin expression in the HOTAIR expressing fibroblasts, but had minimal effect on α-SMA, CTGF, NID and β-catenin expression in the scramble fibroblasts (Figure 1H). Interestingly, at the concentrations analysed, AQB was superior to GSK126 in suppressing HOTAIR mediated fibroblast activation (Figure 1I). Similar results were observed when measuring *ACTA2* transcript levels (Figure 1J). At the tested concentrations, AQB was more potent than GSK126 at reducing HOTAIR mediated *ACTA2* transcription.

### Inhibition of HOTAIR blocks pro-fibrotic gene expression in SSc dermal fibroblasts *in vitro*

Since the AQB anti-fibrotic effects are dependent on HOTAIR overexpression we next sought to assess the effects in SSc dermal fibroblasts. In this context, treatment with AQB at 20µM suppressed the elevated expression of α-SMA, CTGF, NID, GLI2 and β-catenin, while these remained unaffected in healthy fibroblasts (Figure 2A). Consistent with what we observed in HOTAIR overexpressing fibroblasts, also in this context pSMAD3 levels were unaffected (Figure 2A). At the transcriptional level, only 48 h treatment with AQB reduced *COL1A2* (Figure 2B) (20% reduction), *CCN2* (Figure 2C) (25% reduction) and *HES1* (Figure 2D) (30% reduction) transcription in SSc fibroblasts. We observed a dose dependent reduction in NID, α-SMA, CTGF and β-catenin levels in SSc fibroblasts where AQB was able to suppress pro-fibrotic gene expression at a concentration of 10µM and above (Figure 2E). Altogether, these data confirms the importance of HOTAIR/EZH2 interaction in the epigenetic reprogramming that leads to SSc fibroblast activation, as previously shown (2), and supports that notion that specifically targeting this interaction with a small molecule inhibitor can reduce pro-fibrotic gene expression n SSc patient fibroblasts without affecting cellular function of normal dermal fibroblasts.

**Figure 2:**
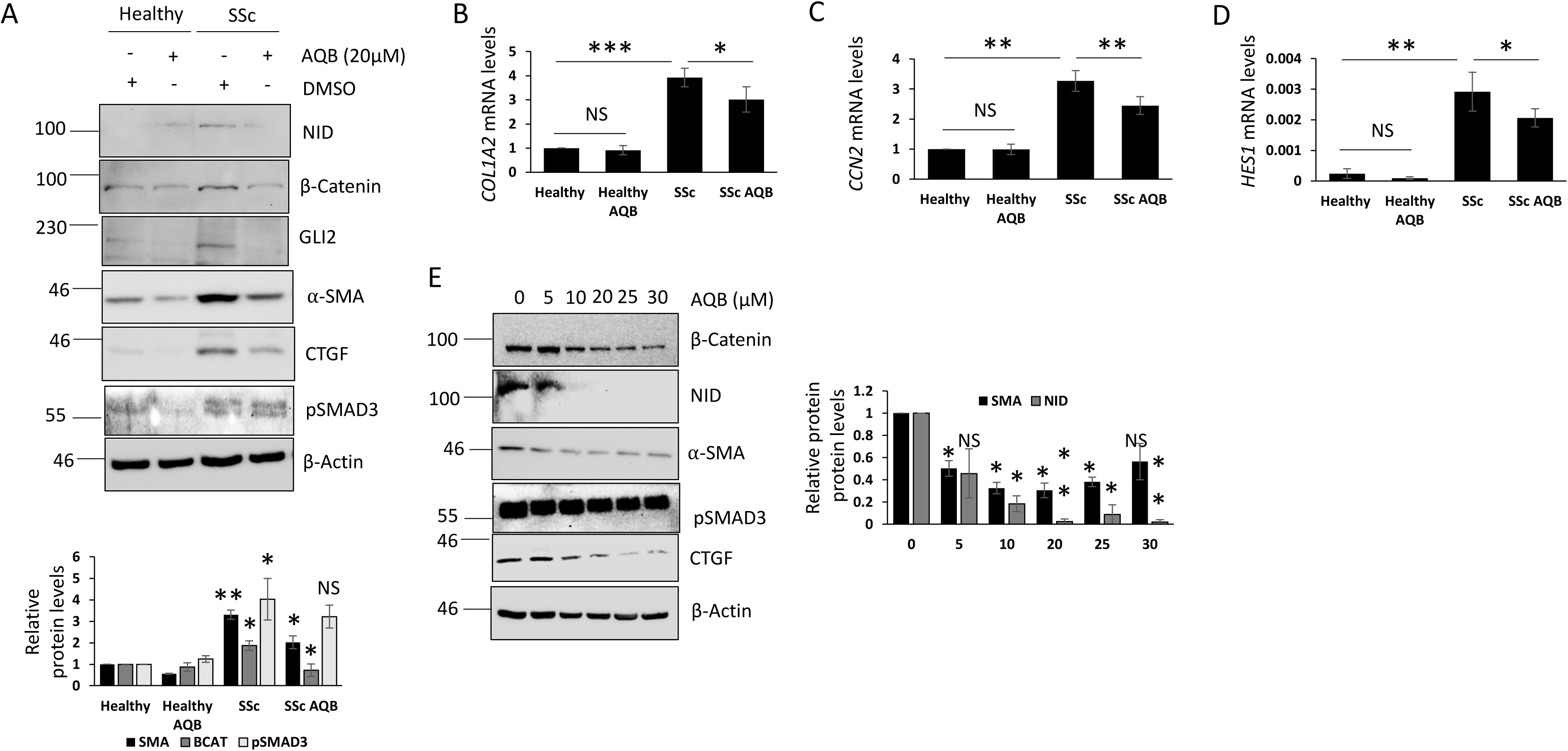
Inhibition of HOTAIR blocks pro-fibrotic gene expression in SSc fibroblasts. Protein and RNA were extracted from healthy and SSc dermal fibroblasts treated with the AQB compound (20μM) for 48 hours. (A) β-catenin, Notch intracellular domain (NID), GLI2, α-SMA, CTGF and pSMAD3 protein levels were assessed by western blot. β-actin was used as a loading control. Graph represents densitometry analysis of Healthy vs SSc fibroblasts plus/minus the inhibitor. Col1a2 (B), CTGF (C) and Hes1 (D) transcript levels were assessed by qPCR. (E) Protein was extracted from SSc fibroblasts treated with a range of concentrations of AQB (5-30µM) for 48 hours. β-catenin, Notch intracellular domain (NID), pSMAD3, CTGF and α-SMA protein levels were assessed by western blot. β-actin was used as a loading control. Graph represents densitometry analysis of SSc fibroblasts treated with the various doses of the inhibitor.

Given the known role of HOTAIR as epigenetic regulator of cellular function, we next set out to investigate if the antifibrotic effect if of AQB was maintained upon drug withdrawal in SSc fibroblasts. For this purpose, SSc dermal fibroblasts were continuously treated with AQB over 4 passages in culture and then cultured for a further 2 passages in the absence of any treatment. Culturing the SSc fibroblasts with AQB for 4 passages strongly reduced α-SMA, CTGF and NID protein levels; however, the levels of NID and α-SMA protein levels returned to DMSO control levels after two passages in the absence of the compound (Figure 3A). Interestingly, CTGF protein levels increased after the interruption of AQB treatment but did not return to the same original level. Similar results were observed at the transcriptional level, where the reduction of *ACTA2* mRNA level returned to pre-treatment levels after interrupting AQB treatment, but not *CTGF* and *COL1A2* mRNAs (Figure 3B). These data support the hypothesis that long term treatment with AQB, may have the potential to re-programme SSc fibroblasts but could require a longer duration of treatment to obtain full re-programming.

**Figure 3:**
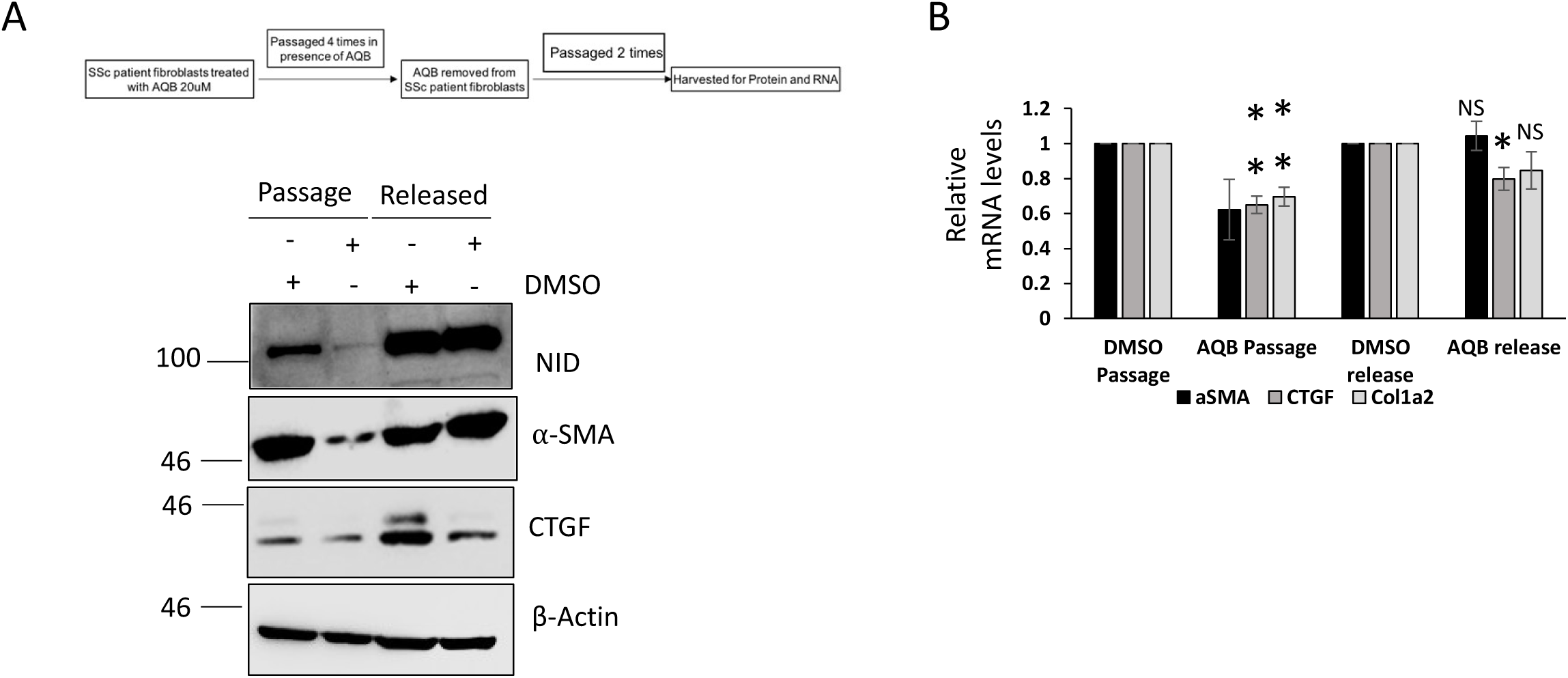
Long term inhibition of the HOTAIR/EZH2 interaction partly re-programmes SSc dermal fibroblasts. SSc dermal fibroblasts were growing in the presence of DMSO or AQB (20μM) for 4 passages and then released from the inhibition and grown for a further 2 passages. Protein and RNA were extracted from both points in the experiment. (A) α-SMA, NID and CTGF protein levels were assessed by western blot. β-actin was used as a loading control. (B) α-SMA, Col1a2 and CTGF mRNA levels were assessed by qPCR.

### Inhibition of HOTAIR/EZH2 interaction prevents paracrine epidermal remodelling

HOTAIR is enriched in the connective tissue of the palmoplantar skin regions (2), areas of skin lacking hair and expressing high levels of Keratin 9 (K9). Affected fibrotic skin areas of SSc patients also typically lack hair suggesting this may be a secondary effect of HOTAIR upregulation. To test this hypothesis we assessed the levels of K9 expression in clinically affected skin from diffuse cutaneous SSc by immunohistochemistry., We found elevated expression of K9 in SSc skin compared to healthy control skin (Figure 4A). This observation informed the hypothesis that similar to palmoplantar skin, the increased HOTAIR expression in SSc dermal fibroblasts may drive induction of K9 in the epidermis of SSc skin. To test this hypothesis, we created organotypic 3D skin equivalent raft cultures containing scramble or HOTAIR expressing dermal fibroblasts underneath a layer of healthy primary dermal keratinocytes (Figure 4B). The epidermis from rafts containing scramble control dermal fibroblasts displayed a typical morphology with a defined cornified layer and showed low expression of K9. on the contrary, the rafts containing HOTAIR-expressing fibroblasts developed an increased epidermal thickness similar to palmoplantar skin, including high K9 expression.

**Figure 4:**
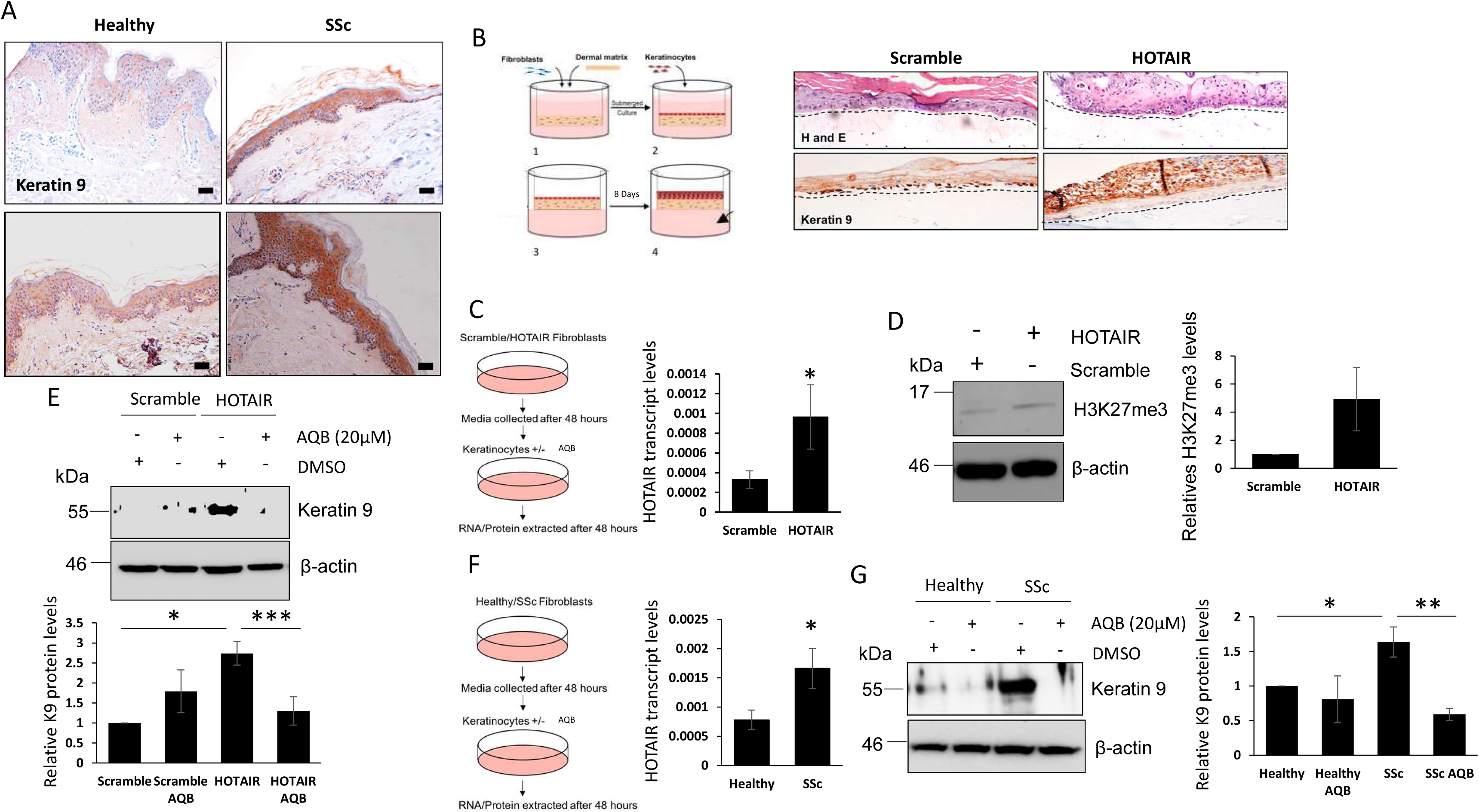
Inhibition of HOTAIR inhibits SSc skin tissue remodelling. (A) Healthy and SSc skin biopsies were stained with an antibody specific to Keratin 9 and counterstained with haematoxylin. Scale bars represent 50μM. (B) 3D skin equivalent skin models were set up with primary keratinocytes and scramble and HOTAIR dermal fibroblasts. Skin sections from the model were stained with H&E and Keratin 9. RNA or Protein was extracted from HaCaTs treated with conditioned media from scramble and HOTAIR expressing fibroblast (C) or healthy and SSc fibroblasts (F). HOTAIR expression levels were assessed by qPCR. (D) H3K27me3 protein levels were assessed in HaCaTs stimulated with scramble and HOTAIR fibroblast conditioned media. HaCaTs were stimulated with conditioned media from scramble and HOTAIR expressing dermal fibroblasts. In addition the cells were treated with the HOTAIR inhibitor AQB. (E) Keratin 9 protein levels were assessed by western blot. β-actin was used as a loading control. Graph represents densitometry analysis of HaCaTs stimulated with scramble and HOTAIR expressing fibroblasts media plus/minus the inhibitor. HaCaTs were stimulated with conditioned media from healthy and SSc dermal fibroblasts. In addition the cells were treated with the HOTAIR inhibitor AQB. (G) Keratin 9 protein levels were assessed by western blot. β-actin was used as a loading control. Graph represents densitometry analysis of HaCaTs stimulated with healthy and SSc fibroblasts media plus/minus the inhibitor

The data above suggest a paracrine signalling between the fibroblasts and keratinocytes which triggers the morphological and transcriptomic changes in the keratinocytes. To explore this possibility, we stimulated the HaCaTs keratinocyte cell line with conditioned media from HOTAIR expressing fibroblasts (Figure 4C) or SSc fibroblasts (Figure 4F). Conditioned media from either HOTAIR-expressing or SSc fibroblasts increased the expression of HOTAIR in the keratinocytes by 2-3-fold compared to the effect of scramble or healthy control fibroblast media (Figure 4C and F). The increase in HOTAIR levels in HaCaTs stimulated with HOTAIR fibroblast conditioned media resulted in a 5-fold increase in H3K27me3 levels compared to scramble fibroblast conditioned media in the HaCaTs cells (Figure 4D). This confirms upregulation of HOTAIR resulted in increased histone methylase activity. These findings are consistent with the previous observation that HOTAIR is increased in both dermis and epidermis in SSc skin sections compared to healthy skin sections as determined by *in situ* hybridization (3). Our *in vitro* skin equivalents data and the observations in SSc patient skin biopsies suggest that the upregulation of HOTAIR in keratinocytes when they are in close proximity to dermal fibroblasts expressing HOTAIR could be necessary for K9 induction in the epidermis. Therefore, we tested if the HOTAIR inhibitor AQB could reverse this phenotype. We stimulated HaCaTs with conditioned media from scramble or HOTAIR-expressing fibroblasts (Figure 4E) and from healthy or SSc fibroblasts (Figure 4G) in the presence of the HOTAIR/EZH2 inhibitor AQB. Conditioned media from HOTAIR and SSc fibroblasts upregulated K9 expression in HaCaTs by 2.7-fold and 1.63-fold, respectively. Interestingly, inhibition of HOTAIR/EZH2 interaction in the keratinocytes prevented the induction of K9 expression by conditioned media from HOTAIR-expressing (Figure 4E) and SSc fibroblasts (Figure 4G). Similar results were observed with the EZH2 inhibitor GSK126. GSK126 treatment blocked the ability of the HOTAIR fibroblast (Supplementary Figure 1A-C) and SSc fibroblast (Supplementary Figure 1D-E) conditioned media from inducing K9 expression in HaCaTs.

### Fibroblasts expressing HOTAIR induce K9 expression and epithelial-to-mesenchymal transition by activation of Wnt signalling

Wnt3a/β-catenin signalling has previously been shown to play an important role in regulating K9 expression in keratinocytes. Treatment of keratinocyte and plantar fibroblast co-cultures with increasing concentrations of a canonical Wnt signalling activator induces *K9* mRNA expression (20). This was reversed when DKK1 (a negative regulator of the Wnt3a/β-catenin signalling pathway) was over-expressed in the keratinocytes (20). Thus, we hypothesised that K9 upregulation in keratinocytes was mediated by Wnt. As previously reported, stimulation of HaCaTs cells with Wnt3a increased expression of K9 at both protein (Figure 5A) and transcript level (Figure 5B). Conditioned media from HOTAIR-expressing and SSc fibroblasts co-upregulated K9 and β-catenin levels (Figure 5C, D), suggesting that Wnt signalling could mediate the effect of HOTAIR on K9 expression. Indeed, inhibition of β-catenin function with FH535 blocked the ability of the HOTAIR (Figure 5C) and SSc fibroblast media to induce K9 expression (Figure 5D). As expected, inhibition of HOTAIR/EZH2 interaction with AQB in the HaCaTs blocked the ability of the HOTAIR (Figure 5E) or SSc fibroblast (Figure 5F) conditioned media to induce β-catenin expression in HaCaTs. Inhibition of the PRC2 complex with GSK126 also prevented SSc fibroblast conditioned media from inducing the expression of β-catenin in keratinocytes (Supplementary Figure 2). Taken together, the data suggests that SSc fibroblast media induces K9 expression in the keratinocytes through a HOTAIR/EZH2/β-catenin signalling cascade.

**Figure 5:**
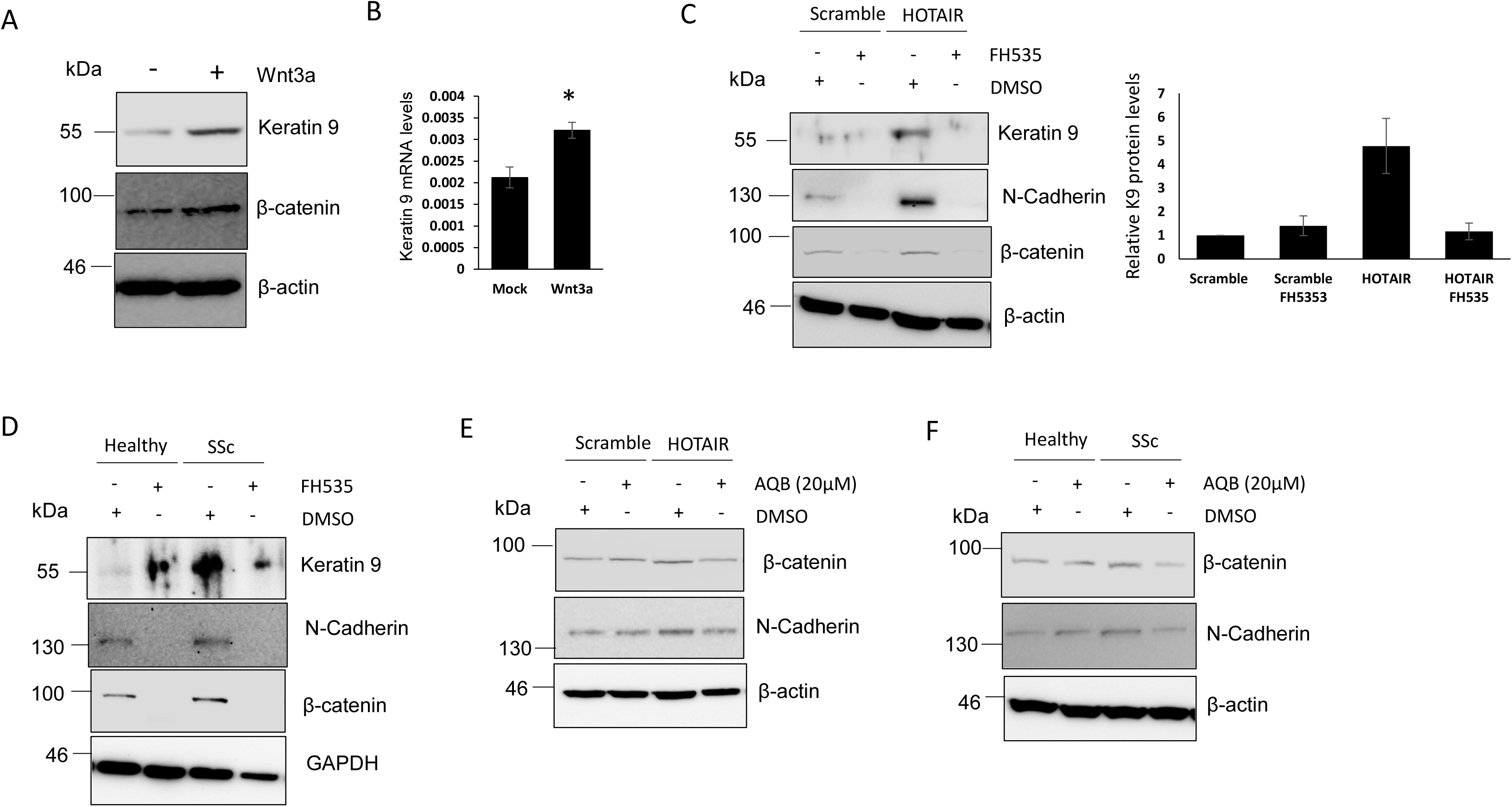
HOTAIR induces Keratin 9 expression in keratinocytes through Wnt3a/β-Catenin signalling. HaCaTs were grown in sera depleted media and stimulated with Wnt3a ligand for 48hrs. RNA and Protein were extracted. (A) Keratin 9 and β-catenin protein levels were assessed by western blot. β-actin was used as a loading control. (B) Keratin 9 transcript levels were assessed by qPCR. HaCaTs were stimulated with conditioned media from scramble and HOTAIR dermal fibroblasts. In addition the cells were treated with the β-catenin inhibitor FH535. (C) Keratin 9, β-Catenin and N-Cadherin protein levels were assessed by western blot. β-actin was used as a loading control. HaCaTs were stimulated with conditioned media from healthy and SSc dermal fibroblasts. In addition the cells were treated with the β-catenin inhibitor FH535. (D) Keratin 9, β-catenin and N-Cadherin protein levels were assessed by western blot. GAPDH was used as a loading control. HaCaTs were stimulated with conditioned media from scramble and HOTAIR expressing dermal fibroblasts. In addition the cells were treated with the HOTAIR inhibitor AQB. (E) β-catenin and N cadherin protein levels were assessed by western blot. β-actin was used as a loading control. HaCaTs were stimulated with conditioned media from healthy and SSc dermal fibroblasts. In addition the cells were treated with the HOTAIR inhibitor AQB. (F) β-catenin and N-Cadherin protein levels were assessed by western blot. β-actin was used as a loading control.

HOTAIR and the Wnt3a/β-catenin signalling pathway been shown to regulate epithelial to mesenchymal transition (EMT) extensively in a range of cancers (21, 22). Keratinocytes in SSc skin have been shown to undergo EMT leading to the transition to myofibroblasts (23). Therefore, we set out to assess whether the HOTAIR mediated induction of β-catenin expression in keratinocytes induced EMT. Indeed, HaCaTs stimulated with HOTAIR fibroblast conditioned media lead to increased expression of N-Cadherin compared to scramble fibroblast conditioned media. This increase in N-cadherin was attenuated when AQB was added (Figure 5E). HaCaTs stimulated with SSc fibroblast media had increased levels of N-cadherin, which was attenuated by AQB (Figure 5F). Further analysis of EMT markers revealed SSc fibroblast media stimulated *TWIST* and *SNAIL* transcription in HaCaTs compared to healthy fibroblast media, while this was attenuated when the HaCaTs were treated with AQB (Supplementary Figure 3)

Similar results were observed in lung epithelial cells (Supplementary Figure 4). A549 cells (Type 1 lung epithelial cell line) were stimulated with scramble and HOTAIR-expressing (Supplementary Figure 4A) or healthy and SSc fibroblast media (Supplementary Figure 4B) with or without AQB. HOTAIR and SSc fibroblast media induced expression of β-catenin, N-cadherin and vimentin in the lung epithelial cells. This induction was attenuated by the HOTAIR/EZH2 inhibitor. This data suggests HOTAIR acting through EZH2 may also play a role in the EMT of lung epithelial cells in SSc.

## Discussion

Targeting lncRNA with small molecule inhibitors is a novel approach. HOTAIR represents a unique therapeutic opportunity as it acts as a scaffold for protein/DNA interactions. Therefore, designing compounds that block the scaffolding functions is feasible. Previous studies have shown the effectiveness of small molecule inhibitors that disrupt HOTAIR binding with EZH2 in breast (8) and glioblastoma (9) cancer models. Here, we show for the first time that these HOTAIR/EZH2 inhibitors block pro-fibrotic gene expression in SSc dermal fibroblasts.

Interestingly, the HOTAIR inhibitor AQB had a similar or higher efficacy than the EZH2 inhibitor GSK126 for blocking pro-fibrotic gene expression in our HOTAIR expressing model system (Figure 1E). This suggests that EZH2 pro-fibrotic functions are entirely dependent on HOTAIR. Therefore, our treatment strategy of directly targeting HOTAIR has merit as we have shown that inhibition of EZH2 is not necessary in the absence of the HOTAIR scaffolding function. This is important as EZH2 inhibitors have been shown to have severe side effects as discussed above.

Long term treatment with AQB partially re-programmed the SSc fibroblasts towards a healthy fibroblast phenotype. Passaging the fibroblasts 4 times in the presence of the inhibitor then releasing the fibroblasts for 2 additional passages, led to a continued suppression of CTGF and COL1A1, suggesting partial re-programming of SSc dermal fibroblasts. Our data suggests that a long-term treatment regime could have anti-fibrotic potential, given its low cytotoxicity.

The ability of SSc dermal fibroblasts to influence keratinocyte biology and functions in a paracrine manner is an emerging field. Our group has shown SSc dermal fibroblasts induce a type I interferon response in keratinocytes through exosome signalling (24, 25) and SSc fibroblast supernatant can activate TGF-β/SMAD signalling in keratinocytes (26). In this study we show that SSc dermal fibroblasts, which express high levels of HOTAIR, induce K9 expression in keratinocytes, both using 3D skin equivalents and using conditioned media. This further highlights the importance of the cross-talk between the cell types in the context of SSc skin. The ability of the SSc fibroblasts to trigger the expression of HOTAIR in the keratinocytes (Figure 3F) may be through the capacity of the fibroblasts to induce TGF-β signalling in the keratinocytes leading to upregulation of HOTAIR (Figure 6), as previous studies have shown TGF-β receptor signalling can trigger HOTAIR transcription in cancer associated fibroblasts (27) Alternatively, HOTAIR may be transported from the SSc fibroblasts to the keratinocytes through exosomes (Figure 6).

**Figure 6:**
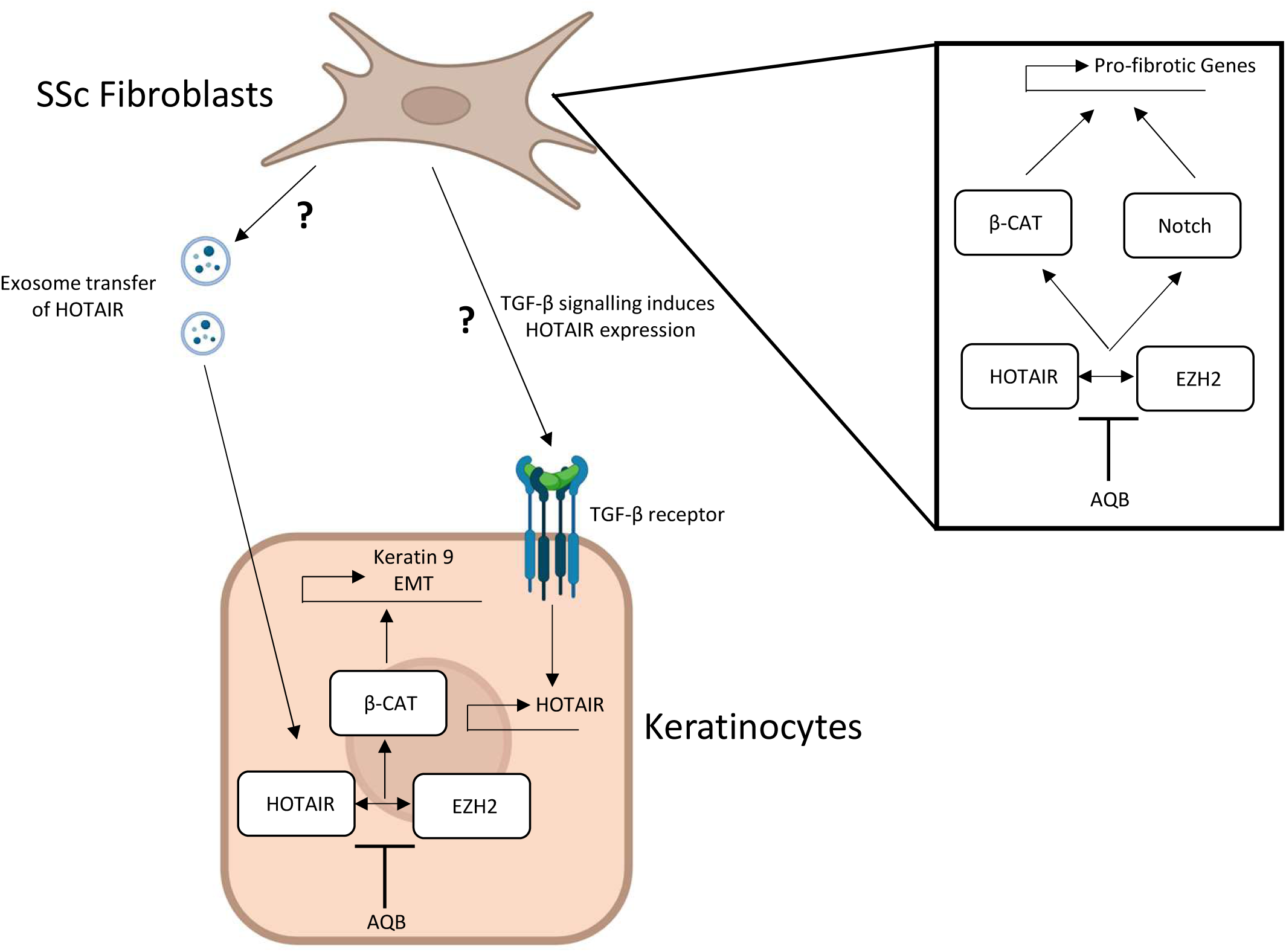
Inhibition of HOTAIR prevents SSc associated tissue re-modelling.

Further analysis revealed SSc dermal fibroblasts can trigger EMT responses in neighbouring keratinocytes and this is in part through HOTAIR as AQB treatment prevented SSc fibroblast conditioned media from inducing N-Cadherin protein expression as well as twist and snail transcription. There is previous evidence that HOTAIR can regulate EMT through its interaction with the co-REST complex and independent of EZH2 (21). Alternatively, HOTAIR in cooperation with EZH2 was shown to be recruited to various EMT related promoters by snail as a means to regulate EMT (28). In the context of SSc and skin keratinocytes we have shown HOTAIR induces EMT in cooperation with EZH2 and this may be driven by the ability of HOTAIR to enhances Wnt signalling in the keratinocytes. Furthermore, it is highly unlikely that HOTAIR regulates EMT in these cells through TGF-β signalling as AQB has been to not alter SMAD3 activation in the context of SSc. It is possible that exosomal transfer of HOTAIR from the fibroblasts to the epithelial cells may drive EMT. Conditioned media from HOTAIR expressing fibroblasts pre-treated with AQB was unable to induce EMT compared to DMSO control media (Figure 4J). Therefore, HOTAIR exported from the fibroblasts maybe associated with the inhibitor and therefore rendered inactive to induce EMT in neighbouring epithelial cells. Overall, the ability of HOTAIR derived from SSc fibroblasts to induce EMT in neighbouring keratinocytes represents a novel origin of myofibroblasts in SSc skin.

Future work will explore the effectiveness of AQB in *ex-vivo* SSc patient skin. SSc skin biopsies will be grown in culture for limited periods of times and pro-fibrotic gene expression will be assessed. Examining the effectiveness of the compounds in mouse skin fibrosis models is not feasible as there is no mouse homologue of HOTAIR.

In conclusion, we have shown for the first time that specific inhibition of HOTAIR/EZH2 interaction with a small molecule inhibitor can block pro-fibrotic gene expression in SSc patient fibroblasts *in vitro* representing a novel therapeutic avenue in the treatment of SSc. Furthermore, the HOTAIR inhibitor has revealed an essential role for HOTAIR in inducing functional changes in the epidermis of skin leading to loss of keratinocyte functions.

## Acknowledgements

**C.W.W** is supported by Susan Cheney Scleroderma fellowship. **F.D.G**, **R.L.R** and **P.Q.M** are supported by the National Institute for Health Research (NIHR) Leeds Biomedical Research Centre (BRC). The views expressed are those of the author and not necessarily those of the NIHR or the Department of Health and Social Care. We would like to thank ChunSheng Kang (Department of Neurosurgery, Tianjin Medical University General Hospital) for kindly providing us with the HOTAIR inhibitor AC1Q3QWB to use in this study.

## Author Contributions

Designing Research Study: C.W.W, N.A.R.D.G, F.D.G,

Conducting Experiments: C.W.W, P.M, S.D, E.B.P, R.L.R

Acquiring Data: C.W.W, P.M, E.B.P, R.L.R

Analysing Data: C.W.W, P.M, N.A.R.D.G, F.D.G

Writing Manuscript: C.W.W, N.A.R.D.G, F.D.G

All authors reviewed and approved the final version of the manuscript

## Ethics Approval and consent to participate

The study was approved by NRES committee NorthEast-Newcastle & North Tyneside: REC Ref:15/NE/0211 to FDG. All participants provided written informed consent to participate in this study. Informed consent procedure was approved by NRES-011NE to FDG by the University of Leeds.

---

**Supplementary Figure 1: HOTAIR induces keratin 9 in keratinocytes through the PRC2 complex.** (A) Conditioned media was collected from scramble and HOTAIR expressing fibroblasts and added to HaCaTs for 48 hours with and without GSK126. (B) K9 protein levels were assessed by western blot. β-actin was used as a loading control. (C) K9 transcript levels were assessed by RTqPCR. (D) Conditioned media was collected from healthy and SSc fibroblasts and added to HaCaTs for 48 hours with and without GSK126. (E) K9 protein levels were assessed by western blot. β-actin was used as a loading control.

**Supplementary Figure 2: HOTAIR regulates Wnt3a/**β**-Catenin signalling through its ability to modulate the PRC2 complex.** Conditioned media was collected from healthy and SSc fibroblasts and added to HaCaTs for 48 hours with and without GSK126. β-catenin protein levels were assessed by western blot. β-actin was used as a loading control.

**Supplementary Figure 3: HOTAIR induces and EMT expression in keratinocytes.** Conditioned media was collected from healthy and SSc fibroblasts and added to HaCaTs for 48 hours with and without AQB. Twist (A) and Snail (B) transcript levels were assessed by RTqPCR.

**Supplementary Figure 4: SSc fibroblasts trigger EMT in lung epithelial through HOTAIR/Wnt3a/**β**-Catenin signalling.** (A) Conditioned media was collected from scramble and HOTAIR fibroblasts and added to A549 cells for 48 hours with and without AQB. N cadherin and β-catenin protein levels were assessed by western blot. β-actin was used as a loading control. (B) Conditioned media was collected from healthy and SSc fibroblasts and added to A549 cells for 48 hours with and without AQB. N cadherin, Vimentin and β-catenin protein levels were assessed by western blot. β-actin was used as a loading control. Graph represents densitometry analysis of A549 cells stimulated with healthy and SSc fibroblasts media plus/minus the inhibitor.

