## Supplementary Figures 1-4 for "The lncRNA HOTAIR/EZH2 interaction inhibitor AC1Q3QWB (AQB) attenuates fibrotic SSc skin tissue re-modelling"

### Slide 1
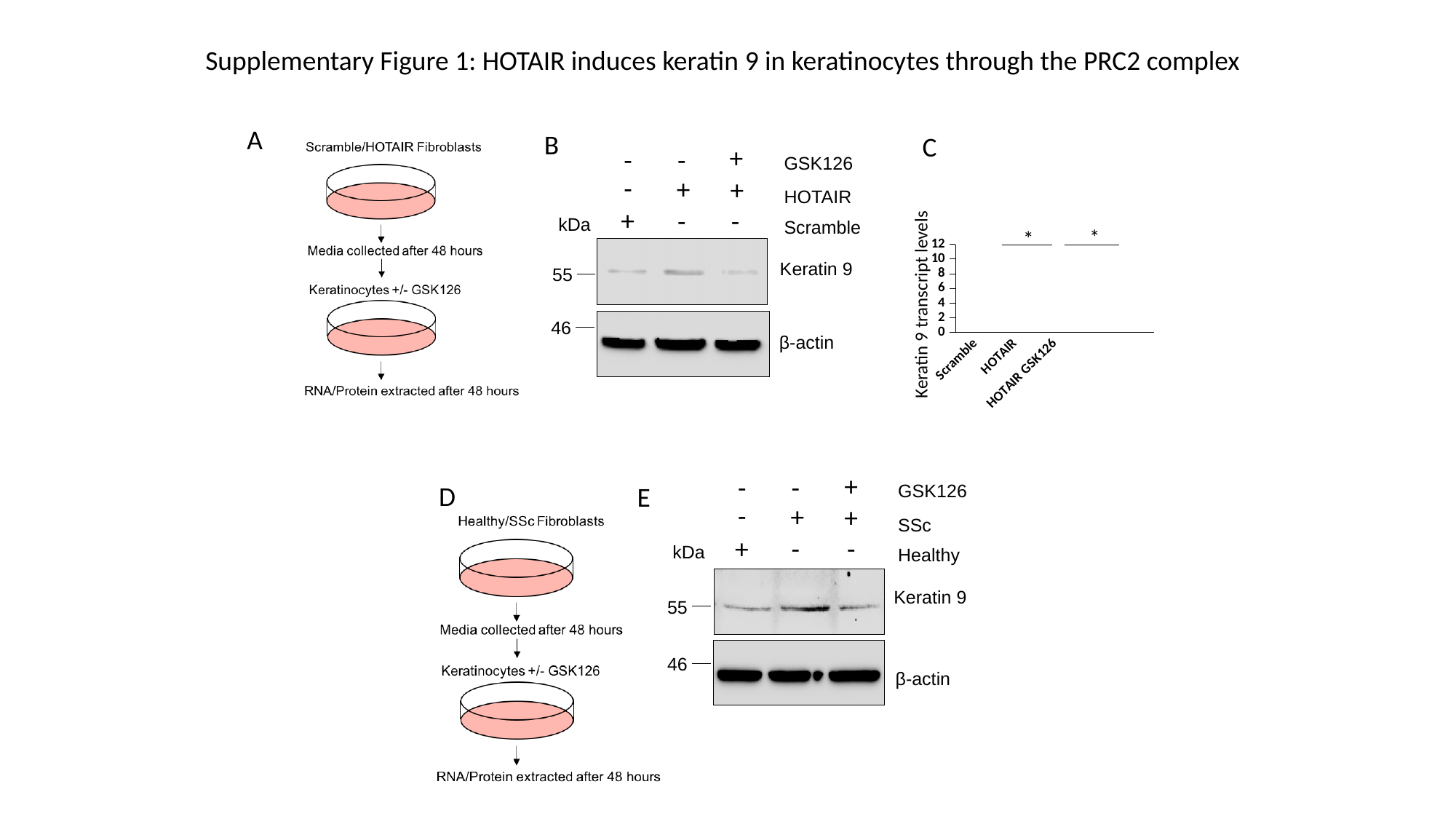

Supplementary Figure 1: HOTAIR induces keratin 9 in keratinocytes through the PRC2 complex
A
B
C
+
-
-
GSK126
-
+
+
HOTAIR
+
-
-
kDa
Scramble
*
*
#### Chart
| Category | ctr |
|---|---|
| Scramble | 0.0021319106 |
| HOTAIR | 0.0039809664 |
| HOTAIR GSK126 | 0.002546256 |
Keratin 9
55
Keratin 9 transcript levels
46
β-actin
+
-
-
D
GSK126
E
-
+
+
SSc
+
-
-
kDa
Healthy
Keratin 9
55
46
β-actin

### Slide 2
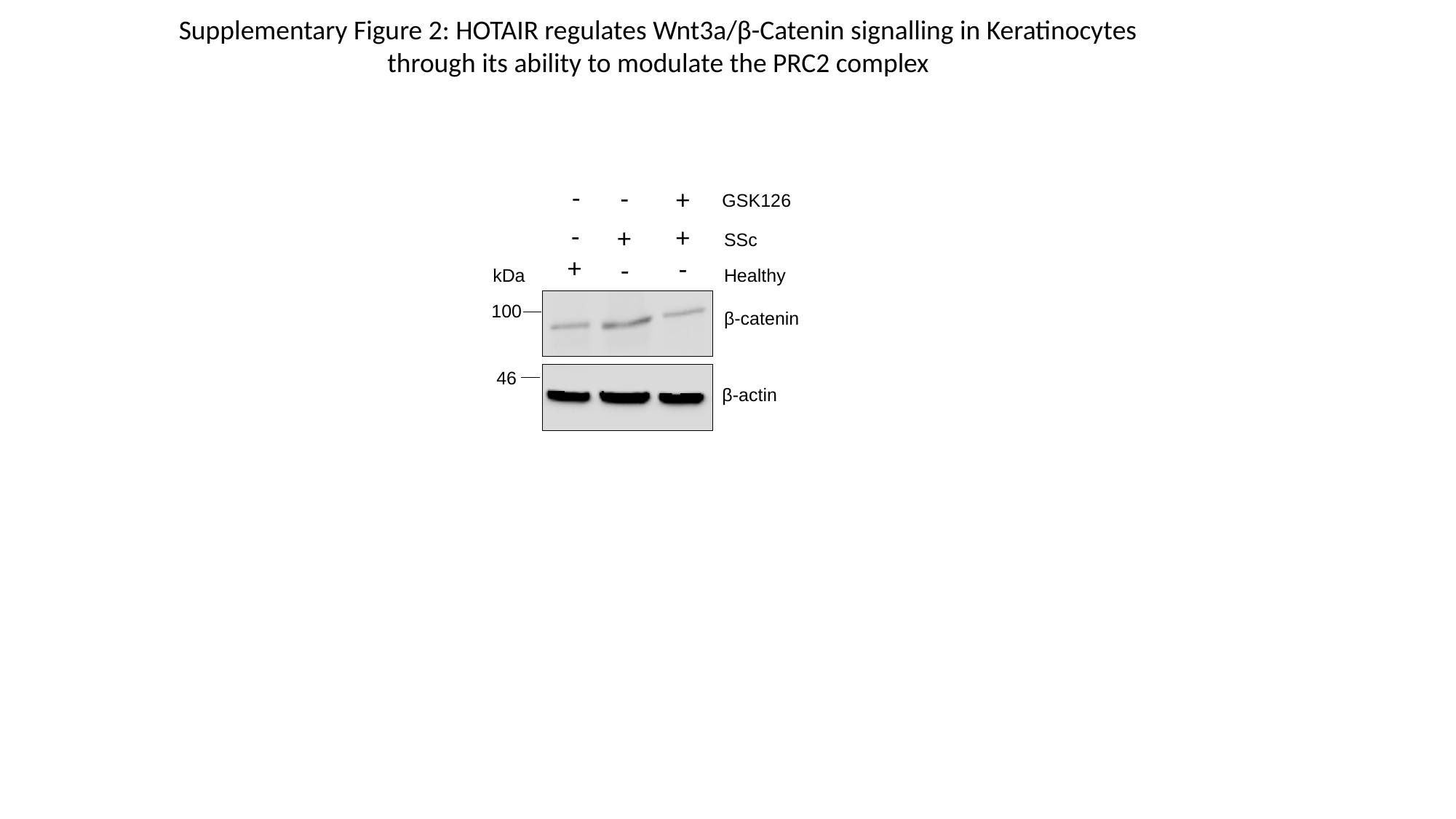

Supplementary Figure 2: HOTAIR regulates Wnt3a/β-Catenin signalling in Keratinocytes
through its ability to modulate the PRC2 complex
-
-
+
GSK126
-
+
+
SSc
+
-
-
Healthy
kDa
100
β-catenin
46
β-actin

### Slide 3
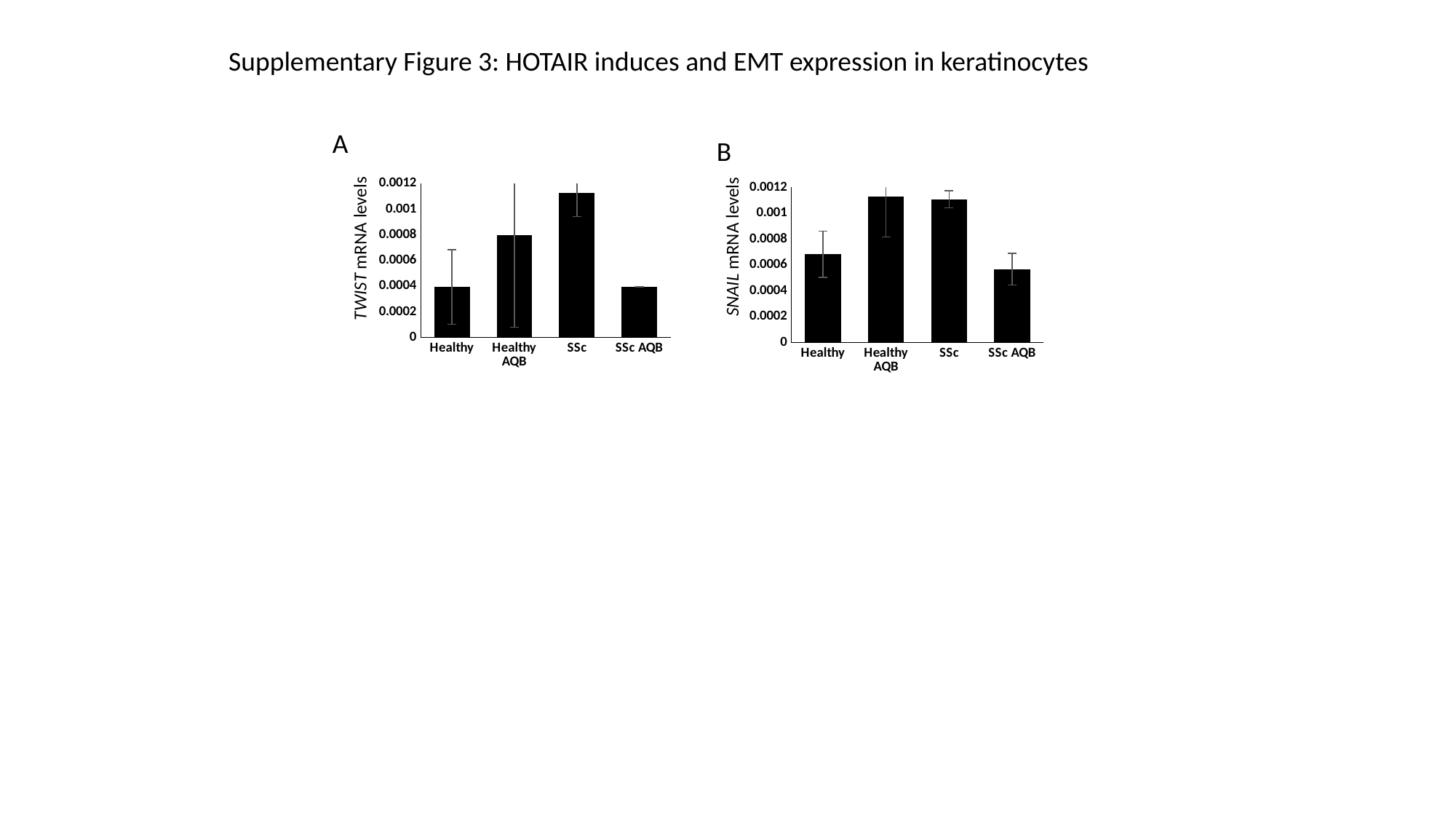

Supplementary Figure 3: HOTAIR induces and EMT expression in keratinocytes
A
B
#### Chart
| Category | Twist |
|---|---|
| Healthy | 0.000392842443667525 |
| Healthy AQB | 0.000800892942901439 |
| SSc | 0.00112947749306178 |
| SSc AQB | 0.000393868046495179 |
#### Chart
| Category | Snail |
|---|---|
| Healthy | 0.000682223057918223 |
| Healthy AQB | 0.00112759278288567 |
| SSc | 0.00110733472978274 |
| SSc AQB | 0.000566762393506173 | SNAIL mRNA levels
 TWIST mRNA levels

### Slide 4
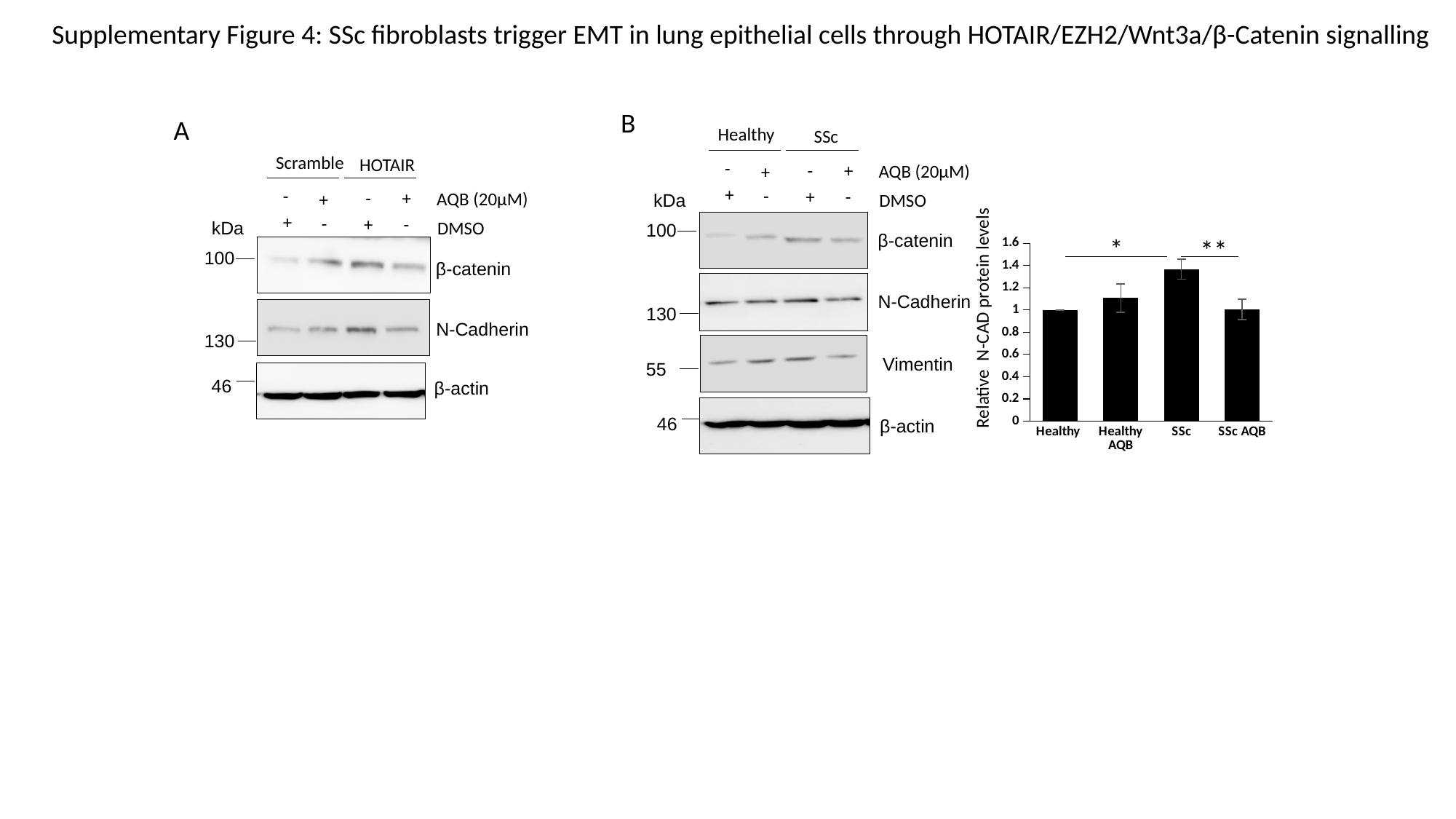

Supplementary Figure 4: SSc fibroblasts trigger EMT in lung epithelial cells through HOTAIR/EZH2/Wnt3a/β-Catenin signalling
B
A
Healthy
SSc
Scramble
HOTAIR
-
-
+
AQB (20µM)
+
+
-
-
+
-
-
+
AQB (20µM)
+
DMSO
kDa
+
-
+
-
DMSO
kDa
100
β-catenin
*
**
#### Chart
| Category | N CAD |
|---|---|
| Healthy | 1.0 |
| Healthy AQB | 1.1075 |
| SSc | 1.36765 |
| SSc AQB | 1.00525 |
100
β-catenin
N-Cadherin
130
Relative N-CAD protein levels
N-Cadherin
130
Vimentin
55
46
β-actin
46
β-actin
